# Machine Learning Reveals Intrinsic Determinants of siRNA Efficacy

**DOI:** 10.1101/2025.08.11.667724

**Authors:** Christian Mandelli, Giulia Crippa, Sathya Jali

## Abstract

Small interfering RNAs (siRNAs) are widely used in therapeutics and agriculture for sequence-specific gene silencing. However, siRNA efficacy remains difficult to predict due to complex dependencies on sequence, structure, and thermodynamic properties. Existing computational tools largely rely on heuristic rules or pre-scored features, limiting generalizability and biological interpretability. Here, we present a machine learning model to predict siRNA efficacy directly from intrinsic antisense sequence features. Using a dataset of 2,428 experimentally validated siRNAs, we developed a comprehensive feature set that encompasses sequence composition, regulatory motifs, thermodynamic parameters, and structural complexity. We trained and evaluated multiple models for both regression and classification tasks. Support Vector Regression (SVR) achieved the best regression performance overall, with a predictive accuracy of R = 0.719 and R^2^ = 0.516, while logistic regression achieved the best classification results with ROC = 0.886 and F1 = 0.809 using a combination of composition, motif, and thermodynamic features. Among all features, position-specific nucleotides were the strongest predictors of efficacy, with a uracil at the 5^′^antisense end (P1_U) and an adenine at the 3^′^end (P19_A) showing the highest influence, consistent with known mechanisms of strand selection and RISC loading. Our approach improves both predictive power and biological interpretability compared to existing methods, eliminating reliance on external scoring functions. The resulting framework supports rational siRNA design for therapeutic applications, functional genomics, and non-transgenic crop protection strategies.

## Introduction

RNA interference (RNAi) is a conserved, sequence-specific gene regulatory mechanism found across eukaryotes, serving essential roles in antiviral defense, epigenetic modulation, and post-transcriptional gene silencing. Central to RNAi are small interfering RNAs (siRNAs), which originate from the cleavage of long double-stranded RNA (dsRNA) by Dicer-like (DCL) enzymes into 21- and 24-nucleotide duplexes. siRNAs are incorporated into Argonaute (AGO) proteins to form the RNA-induced silencing complex (RISC), which guides mRNA degradation or transcriptional silencing via RNA-directed DNA methylation (RdDM) in plants or translational repression in animals (1, 2). These processes are not only fundamental to eukaryotic gene regulation but also represent powerful entry points for targeted genetic intervention.

Initially developed in model systems, siRNAs have become a clinically validated therapeutic platform, with several approved drugs—such as patisiran for hereditary transthyretin amyloidosis (3)—and expanding pipelines targeting previously inaccessible genes (2). By acting at the mRNA level, siRNAs provide a mechanism different from traditional small molecules or antibodies, enabling precise, transient, and programmable gene knockdown. In agriculture, RNAi technologies are used to suppress pathogens and modulate plant traits through methods such as host-induced gene silencing (HIGS) and virus-induced gene silencing (VIGS). More recently, spray-induced gene silencing (SIGS) has emerged as a promising non-transgenic approach, allowing topical application of double-stranded RNA (dsRNA) to provide protection without genetically modifying the host plant. These applications demonstrate the broad potential of RNAi translation—yet they also reveal a key challenge: siRNA design still relies on an empirical, unpredictable process.

A major obstacle to effectively using siRNA-based technologies—whether for therapy or agriculture—is the inconsistent effectiveness of siRNA. Although sequence complementarity is necessary for target binding, it does not ensure successful silencing. DCL processing of dsRNA can produce diverse siRNA pools with different strengths, and small changes in siRNA sequences can influence strand selection by RISC, leading to lower effectiveness or unintended off-target effects (4, 5). These effects reflect deeply embedded biophysical and biochemical limitations that affect RNAi efficiency at various stages, from initial duplex processing to target access and AGO loading. Additionally, siRNA activity is affected by a complex interplay of sequence, thermodynamics, structure, and context, many of which are difficult to predict with traditional heuristic or rule-based design methods. As a result, designing effective siRNAs remains a lengthy, error-prone process, which limits wider usage and scaling of RNAi in both medicine and agriculture.

Several computational tools for siRNA design have been developed—including siDirect, i-Score, RNAxs, Biopredsi, dsCheck, and si-Fi—which largely rely on empirically derived design rules or linear models, emphasizing sequence complementarity, GC content, or thermodynamic asymmetry (6–10). These tools have provided valuable guidance, particularly in early development, but as highlighted in (2), they often fail to account for the complex biology behind RNAi. Notably missing from many platforms are considerations of mRNA target accessibility, local RNA secondary structure, siRNA duplex stability, and off-target interactions—all of which influence silencing efficiency. Without incorporating these biological factors, such tools fail to perform well across different organisms, cell types, or experimental conditions (9, 11–13).

A significant breakthrough was achieved with the *pssRNAit* platform (5), which integrates multiple siRNA prediction tools within a cascade support vector machine (SVM) framework. More recently, deep learning–based methods such as *DeepSilencer* (14) have further advanced the field by leveraging transformer–CNN hybrid architectures and multi-task learning, achieving state-of-the-art performance for siRNA knockdown prediction. However, these approaches often require large datasets and are less interpretable, limiting their applicability, thus rational siRNA design.

In this work, we present a machine learning framework that significantly enhances siRNA efficacy prediction by only leveraging intrinsic 21 nt antisense siRNA sequence features, eliminating reliance on external scoring systems. Our approach integrates rigorous feature engineering—encompassing sequence composition, regulatory motifs, thermodynamic properties (e.g., ΔG, melting temperature), and structural complexity from RNAfold—with advanced nonlinear models, including Support Vector Regression (SVR) and XGBoost. Our results demonstrate high predictive accuracy for both regression (*R* = 0.719, *R*^2^ = 0.516) and classification tasks (*ROC* = 0.886, *F*1 = 0.809) using a combination of composition, motif, and structural features, outperforming linear models and single-feature approaches, while maintaining full biological interpretability and computational efficiency.

## Materials and Methods

Data and codes are available on Github.

### Dataset

To ensure comparability with existing studies, we used the well-established siRNA efficacy dataset from (15). In detail, our dataset comprises 2,428 experimentally tested siRNAs targeting 34 mRNAs, derived from a reporter gene assay and employed in training the BIOPREDsi model.

For classification-based evaluations, two efficacy thresholds — 0.5 and 0.7 — were applied to convert continuous efficacy scores into binary labels for high versus low activity.

### Features Extraction

To enable predictive modeling of siRNA efficacy, a custom Python-based feature extraction pipeline that processes each 21-nucleotide antisense siRNA strand and encodes it into a structured numerical format suitable for our methodology was developed. A description of all features can be found in Table SN.1.

Recent work has shown that distinct classes of sequence-derived features contribute differentially to siRNA efficacy, with some influencing strand selection, others affecting target accessibility, and still others modulating off-target risk (2). Therefore, features from four primary categories were extracted: (1) sequence composition, (2) regulatory and structural motifs, (3) thermodynamic parameters, and (4) structural complexity.

The composition-based features include both position-specific nucleotide identities and global nucleotide distributions. These capture local sequence preferences known to influence strand selection and siRNA functionality (15). Global composition metrics such as GC content and skew between the 5^′^ and 3^′^ ends were included due to their relevance to thermodynamic asymmetry and RISC incorporation (16, 17).

To account for regulatory and structural motifs, patterns at both termini of the siRNA duplex were identified, including recurring sequence motifs, palindromic structures, and simple nucleotide repeats. These elements have been previously associated with enhanced silencing efficiency, reduced off-target potential, or increased structural instability (4, 18–20).

Thermodynamic features were computed to capture both strand and duplex stability. This includes the free energy of hybridization with the target, the overall folding energy of the antisense strand, melting temperature estimates, and nearest-neighbor stacking energies. Local asymmetries across specific regions of the duplex were also evaluated using sliding-window calculations of free energy differences (18, 21–23).

Lastly, structural complexity features from RNA secondary structure predictions were generated using RNAfold (version 2.6.4) from the ViennaRNA Package (24). These include metrics related to folding energy, base-pairing density, loop topology, and nucleotide-level entropy. These properties inform on the accessibility and rigidity of the siRNA structure, both of which impact RISC loading and functional efficiency (2, 25).

All categorical features were one-hot encoded to ensure compatibility with numerical values. The resulting feature matrix was exported as a CSV file and used as input for downstream model training and testing, for both regression and classification tasks.

### Dataset Preprocessing and Analysis

Prior to model training, standard cleaning procedures were applied to ensure data integrity and consistency. All infinite values were replaced with missing values and subsequently dropped. A small number of sequences exhibiting extreme global folding energy (Global Δ*G* > 100 or < − 100 kcal/mol) were removed, as these likely represent non-biological outliers or computational artifacts from RNA secondary structure prediction. This reduced the dataset from 2,428 to 2,408 siRNAs.

All numerical features were normalized, and categorical variables were one-hot encoded. The final preprocessed dataset contained 189 features, including one-hot encodings of positional nucleotides and selected di- and trinucleotide motifs. The structure of key thermodynamic and compositional features after preprocessing is shown in Figure SN.1, which illustrates distinct pairwise correlations and unimodal distributions, supporting the biological relevance and diversity of the cleaned dataset. Lastly, to address class imbalance in the binary classification task, a label balancing procedure was applied prior to training. The dataset was re-sampled using undersampling. Specifically, an equal number of positive and negative labels was randomly selected to form a balanced training set.

### Machine Learning Model Training and Evaluation

To assess the predictive power of the extracted siRNA features, a comprehensive machine learning framework encompassing both regression and classification tasks was implemented. All models were trained and evaluated using a unified pipeline based on scikit-learn, ensuring consistency in preprocessing, feature transformation, and validation procedures. Prior to model fitting, the full dataset was randomly split into training (80%) and testing (20%) subsets. For classification tasks, the split was stratified by class label to preserve class proportions. Continuous features were standardized using StandardScaler, while binary (categorical) features were passed through without transformation.

For regression analysis, the target variable was treated as continuous, and three predictive models were evaluated: ordinary least squares linear regression (LR), support vector regression (SVR) with a radial basis function kernel, and gradient-boosted regression trees (XGBoost). Model performance was assessed using 5-fold cross-validation repeated across all training data partitions. The evaluation metrics included mean absolute error (MAE), root mean squared error (RMSE), coefficient of determination (*R*^2^), and Pearson’s correlation coefficient (*R*). In addition to the cross-validation metrics, each model was also evaluated on the held-out test set to assess its generalization capacity.

To investigate the contribution of different feature classes to model performance, models were systematically trained on all 2^*k*^− 1 non-empty combinations of *k* = 4 biologically motivated feature categories: sequence composition, regulatory motifs, thermodynamic stability, and structural complexity. For each feature subset, the corresponding columns were extracted from the training and test data, processed independently, and passed to the model pipeline. This exhaustive combinatorial design enabled direct quantification of the marginal and joint predictive utility of each biological feature group across both regression and classification objectives.

A parallel classification pipeline was implemented to predict binary siRNA efficacy labels (effective vs. ineffective). For this task, four classifiers were trained: logistic regression (LR), support vector machines (SVM), random forests (RF), and multi-layer perceptron neural networks (NN). As in the regression setting, models were trained using 5-fold stratified cross-validation, with performance evaluated using accuracy, precision, recall, F1 score, and the area under the receiver operating characteristic curve (ROC). Class imbalance was mitigated using undersampling, ensuring an equal representation of positive and negative examples in the training set. Test performance was computed separately on the 20% hold-out set.

All models were implemented using standard Python libraries, including scikit-learn for linear models, SVMs, and neural networks, and XGBoost for gradient-boosted regression. The evaluation pipeline and model training loops were designed to support modular experimentation across feature sets, model types, and target modalities. This framework enabled a rigorous comparison of both regression and classification strategies and allowed us to quantify how different combinations of siRNA features influence predictive accuracy.

### Computational Pipeline

To ensure full reproducibility, all analyses described above were implemented as an automated Nextflow (DSL2) pipeline (26). The pipeline takes a single FASTA file as input and executes the complete workflow from feature extraction through model training, evaluation, and generation of all figures and tables reported in this study without manual intervention. The pipeline comprises 18 modular processes organized into five stages: (1) feature extraction and train/test splitting, (2) regression and classification model training, (3) systematic feature group evaluation, (4) permutation importance analysis, and (5) figure and table generation. Processes without data dependencies execute in parallel, while dependent stages (e.g., Table 1 synthesis from supplementary tables) are automatically coordinated by Nextflow’s dataflow model. All random seeds are fixed (`random_state=42`) throughout, and dependencies are managed via a Conda environment specification. The complete source code is available at https://github.com/sathyasjali/siRNA_ML.

**Table 1.** Results summary (test set performance). Models trained on different feature group combinations, sorted by *R* for regression and ROC AUC for classification. Best values are highlighted. C: Composition features; M: Motif and regulatory elements; T: Thermodynamic stability features; S: Structural complexity features.

| Task | Model | Feature Set | MAE | $R^2$ | $R$ |
| --- | --- | --- | --- | --- | --- |
| Regression | SVR | C + M + S | 0.106 | **0.516** | **0.719** |
|  | SVR | C + M + T + S | 0.106 | 0.515 | 0.719 |
|  | SVR | C + M + T | 0.107 | 0.513 | 0.718 |
|  | XGBoost | C | 0.110 | 0.473 | 0.689 |
|  | LR | C + M + T + S | 0.112 | 0.454 | 0.676 |
| Task | Model | Feature Set | Accuracy | F1 | ROC |
| Classification<br>*Label threshold: 0.5* | Logit | C + M + T | 0.801 | 0.809 | **0.885** |
|  | Logit | C + M | 0.801 | 0.809 | 0.885 |
|  | Logit | C + M + T + S | 0.795 | 0.809 | 0.882 |
|  | RF | C + T | 0.789 | 0.791 | 0.852 |
|  | SVM | C + M | 0.708 | 0.713 | 0.806 |
| Classification<br>*Label threshold: 0.7* | SVM | C + S | 0.760 | 0.763 | **0.844** |
|  | SVM | C | 0.754 | 0.759 | 0.844 |
|  | SVM | C + T | 0.754 | 0.759 | 0.843 |
|  | Logit | C + M + T + S | 0.740 | 0.740 | 0.839 |
|  | RF | C + M | 0.765 | 0.761 | 0.821 |

## Results

In this section, we discuss the results of the selected machine learning models for siRNA efficacy using both regression and classification tasks. Models were trained on combinations of four biologically feature categories: Composition (C), Motifs (M), Thermodynamic (T), and Structural (S) features.

### Nonlinear Models and Composition Features Drive Superior siRNA Efficacy Prediction

Regression models were trained to predict continuous efficacy scores. Among the three tested algorithms – Linear Regression (LR), Support Vector Regression (SVR), and XGBoost – nonlinear methods substantially outperformed linear ones. Table 1 summarizes the test set performance of the top-performing models and feature combinations.

Among all algorithms, SVR achieved the highest performance (*R* = 0.719, *R*^2^ = 0.516) using composition, motif, and structural feature (C+M+S). XGBoost and LR models performed slightly below SVR, particularly when structural and thermodynamic features were used alone.

Across all methods, models trained on composition features (C) consistently outperformed those trained on motif (M), thermodynamic (T), or structural (S) features alone. For instance, in the XGBoost model, using only composition features outperforms (*R* = 0.689, *R*^2^ = 0.743) all other features composition.

Linear models achieved slightly lower predictive performance overall. The best linear model required all features (*R* = 0.676, *R*^2^ = 0.451), underscoring the utility of non-linear learners for capturing complex interactions between siRNA sequence and functions.

### Composition-Based Features Enable Interpretable Models to Outperform Deep Learning in siRNA Efficacy Classification

We evaluated three classifiers – Logistic Regression (LR), Support Vector Machines (SVM), and Random Forests (RF) – to predict binary siRNA efficacy at two different thresholds levels (0.5 and 0.7). Performance was benchmarked using test F1-score and ROC.

As shown in Table 1, logistic regression models trained on composition, motif, and thermodynamic features (C+M+T) yielded the best performance at the 0.5 threshold (ROC = 0.886; F1 = 0.809). On the other hand, SVM trained on composition and structure (C+S) was the top performer at the 0.7 threshold (ROC = 0.844; F1 = 0.763). As per the regression results, features sets lacking composition showed substantially reduced accuracy (ROC 0.73), confirming that sequence composition remains the dominant predictor of efficacy.

Compared to the recently proposed deep learning model DeepSilencer (AUC = 0.820, F1 = 0.775 in cross-dataset evaluation) (14), our logistic regression model achieved higher AUROC (0.886) and F1 (0.809) on the Huesken dataset. Although direct comparison is limited by differences in evaluation protocol (within-dataset vs. cross-dataset), these results suggest that a feature-engineered and interpretable model can achieve competitive or superior performance to deep learning approaches on this benchmark dataset while remaining computationally efficient and biologically transparent.

### Position-Specific Nucleotides and Local Motifs Drive siRNA Efficacy

Feature importance analysis of the best-performing regression model (SVR trained on composition, motif, and structural features; C+M+S) revealed that position-specific nucleotides are the strongest predictors of siRNA silencing efficiency (Figure 1). The most informative feature was a uracil at the 5^′^ antisense end (P1_U), followed by an adenine at the 3^′^ antisense end (P19_A). Other important terminal positions included guanine at position 1 (P1_G) and cytosine or guanine at position 19 (P19_C, P19_G). Short sequence motifs also contributed to prediction accuracy: the UCG trinucleotide motif (UCG%) and the UG dinucleotide (UG%) frequencies ranked within the top 10 features, suggesting that local nucleotide context beyond single positions can modulate siRNA activity. Structural and compositional features such as global GC content (GC_Content) and GC skew between the 5^′^ and 3^′^ ends (GC_skew_5p_vs_3p) had moderate predictive value, while AU content at the 5^′^ end (five_prime_AU) was among the lower but still relevant contributors. Overall, these results indicate that effective siRNAs are primarily defined by specific terminal base identities (especially P1_U and P19_A) together with favorable short sequence motifs and moderate GC-related features, rather than by global thermodynamic properties alone.

**Fig. 1.**
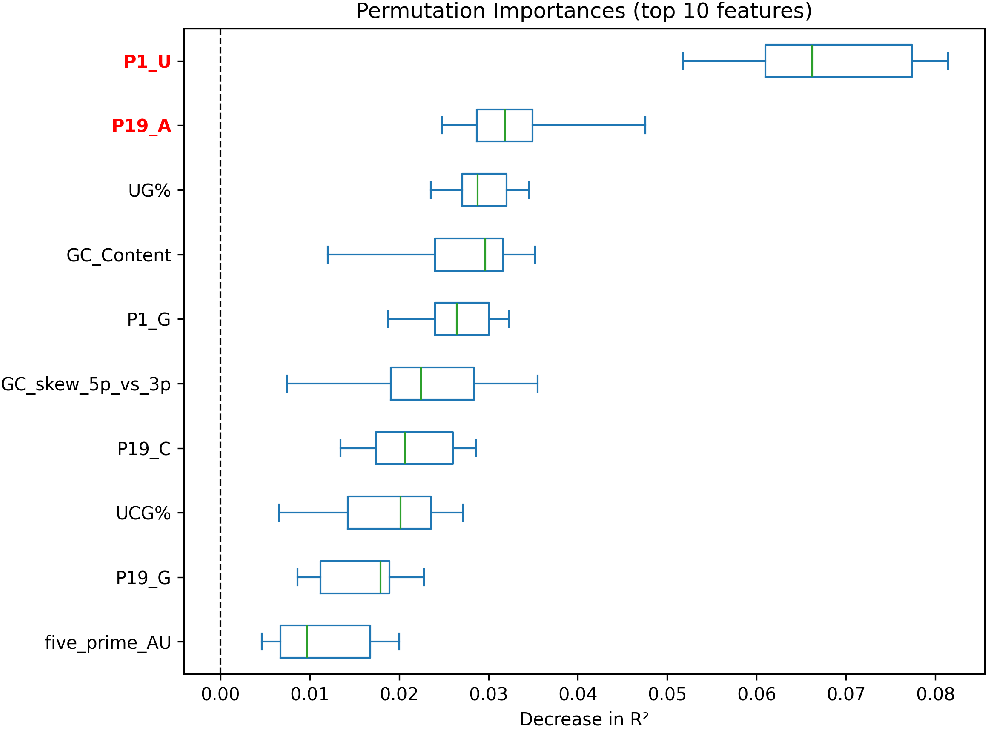
Top 10 most important features ranked by permutation importance from the best-performing regression model: SVR trained on composition, motif, and structural features (C+M+S). Error bars represent standard deviation across 10 shuffled folds. Results from Ridge Regression, XGBoost, and classification tasks show consistent patterns, highlighting the robustness of position-specific features such as P1 and P19.

## Discussion

Our machine learning framework for predicting siRNA efficacy represents a significant advancement over existing computational tools, achieving high performance in both regression and classification tasks. For continuous efficacy prediction, Support Vector Regression (SVR) trained on composition, motif, and structural features (C+M+S) the best performance (*R* = 0.719, *R*^2^ = 0.516). For binary classification, logistic regression trained on composition, motif, and thermodynamic features (C+M+T) outperformed competing models (*ROC* = 0.886, *F*1 = 0.809).

Across both regression and classification tasks, sequence composition features emerged as the most informative and biologically meaningful predictors (1). Notably, P1_U (uracil at the 5^′^ end of the antisense strand) and P19_A (adenine at the 3^′^ end) were the top contributors to model performance, as evidenced by their consistently high permutation importance across 10 random shuffles in our best-performing regression model (Figure 1).

Importantly, this pattern was not unique to SVR: analyses of other regression and classification models (Appendix Figures SN.2 and SN.3) revealed a strikingly similar ranking of features, with P1_U and P19_A repeatedly among the top predictors regardless of the algorithm^1^. Additional terminal nucleotides, such as P19_C and P1_G, also ranked highly in several models, further supporting the functional significance of both ends of the antisense strand.

The importance of P1_U aligns with well-established RNAi principles. Thermodynamic asymmetry at siRNA duplex ends facilitates strand selection by RISC. A uracil at the 5^′^ antisense end decreases local duplex stability, thereby promoting antisense strand loading. This mechanism is supported by prior studies showing that a less stable 5^′^ antisense end—enriched in U or A bases—favors guide strand selection (16, 27). Similarly, (28) demonstrated that Dicer discriminates effective siRNAs by sensing thermodynamic properties of the terminal base pairs, underscoring the functional relevance of position 1 nucleotides.

By contrast, P19_A highlights a less-characterized but potentially important determinant of siRNA efficacy. Our model suggests that adenine at the 3^′^ of the antisense strand enhances RISC loading, likely by influencing how the 3^′^ over-hang is presented to the PAZ domain. This finding is consistent with (29), who showed that optimal PAZ–overhang interactions promote siRNA potency and specificity. The strong predictive contribution of P19_A suggests that 3^′^-end composition plays a more active role in RISC engagement than previously appreciated. Consistent with this, other nucleotides at position 19, including cytosine (P19_C) and guanine (P19_G), also showed high importance (Figure 1), further supporting a key role of terminal base identity in siRNA efficacy.

While, global GC content ranked among top predictors ( ≈ 0.03 importance), its contribution was modest compared to P1_U, position-specific sequence terminal nucleotides can be more informative than aggregate composition measures for accurate efficacy prediction. Beyond composition, regulatory motifs such as the UCG trinucleotide motif (UCG%) and the UG dinucleotide frequency (UG%) were also ranked among the top predictors. These motifs may enhance RISC loading or siRNA–mRNA pairing, though experimental evidence remains limited. Prior studies have noted that such motifs can affect both immunogenicity and off-target effects (30, 31), suggesting a dual role in efficacy and safety.

In addition to sequence motifs, several composition-derived metrics, including GC_skew_5p_vs_3p and five_prime_AU, contributed moderately to prediction accuracy. These features likely capture differences in end stability that facilitate strand selection and guide strand incorporation.

Although individual thermodynamic features such as global melting temperature (Global_Tm) ranked highly in permutation importance analyses, others such as duplex free energy (Δ*G*) did not. Nevertheless, the best-performing regression model (C+M+S) excluded the entire thermodynamic category without any meaningful loss in accuracy (*R*^2^ = 0.516 vs. 0.515 for C+M+T+S). This indicates that thermodynamic information is largely redundant and effectively captured by composition features, particularly GC content, which strongly correlates with duplex stability (27). In contrast, structural features derived from RNAfold predictions— despite modest individual importance—provided significant additive value as a group. Models incorporating these structural features (C+M+S) outperformed those relying on thermodynamic alternatives (C+M+T), likely because they capture local base-pairing density and target site accessibility, which critically influence guide strand unwinding, RISC incorporation, and hybridization efficiency (29). Overall, effective siRNAs appear to strike a balance between structural rigidity (for stable loading into RISC) and sufficient flexibility/local accessibility (for target engagement), whereas global thermodynamic metrics lack the positional specificity and generalizability needed for robust prediction.

Overall, our model advances both predictive accuracy and biological interpretability. The consistent importance of P1_U and P19_A, together with informative short motifs and compositional metrics such as UCG% and GC_skew, highlights their potential as rational design principles for siRNA optimization. Their prevalence in effective siRNAs across a dataset of 2,428 siRNAs suggests broad generalizability.

Our findings have practical relevance for both therapeutic and agricultural RNAi applications. In medicine, they provide a biologically grounded framework for rational siRNA drug design, potentially reducing the need for extensive experimental screening. In agriculture, they can guide SIGS strategies, enabling non-transgenic crop protection (32). The interpretability of our feature set also facilitates mechanistic insights into siRNA function, which can improve understanding of RNAi dynamics.

Despite these advances, several limitations remain. Our training data were derived primarily from in vitro reporter systems, which may not fully capture the complexity of endogenous transcript environments. The strong influence of sequence composition might also reflect dataset-specific biases. Future work should focus on (i) validation across diverse biological systems, including primary cells and whole plants; (ii) integration of higher-order transcriptomic features, such as RNA-binding protein occupancy and epitranscriptomic modifications (m^6^A-seq), which are known to affect RNA folding, stability, and accessibility (33, 34); and (iii) iterative refinement with experimental feedback, ensuring that computational predictions translate effectively to real-world applications.

Finally, we propose that siRNA efficacy reflects a multilayered code integrating sequence, structure, and dynamic transcriptomic context. Achieving higher predictive power will likely require models that combine data-driven learning with biophysical principles. Physics-informed machine learning (PIML) (35, 36) offers a promising approach by embedding mechanistic constraints (e.g., RNA thermodynamics and RISC assembly rules) directly into statistical models. Such hybrid frameworks could improve generalization across biological systems while retaining mechanistic interpretability, bridging the current gap between computational performance and biological validity.

In summary, our study (i) identifies P1_U and P19_A as the most robust and biologically meaningful predictors of siRNA efficacy, (ii) demonstrates that the most predictive individual feature (P1_U) is position-specific rather than a global metric, and that terminal nucleotide identity collectively provides information beyond what GC content alone captures, (iii) provides a biologically interpretable model that can guide rational siRNA design for therapeutic and agricultural applications, and (iv) outlines a path toward hybrid biophysical– statistical models that enable context-aware and mechanistically grounded RNAi prediction.

## Supplementary Note 1: Dataset

**Table SN.1.**
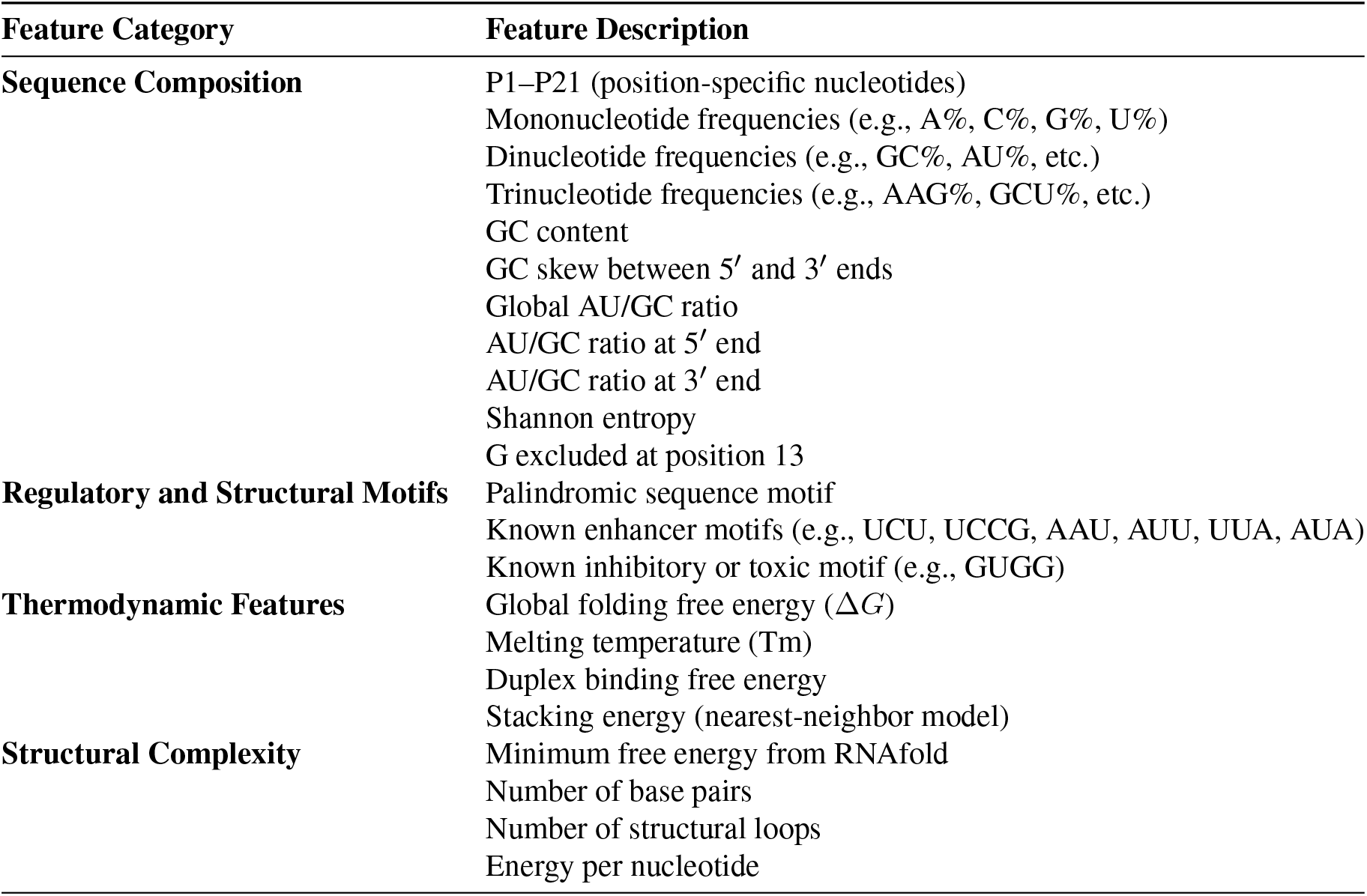
Summary of features used for siRNA efficacy modeling, grouped by category. Full details and abbreviations are provided in the Methods section.

**Fig. SN.1.**
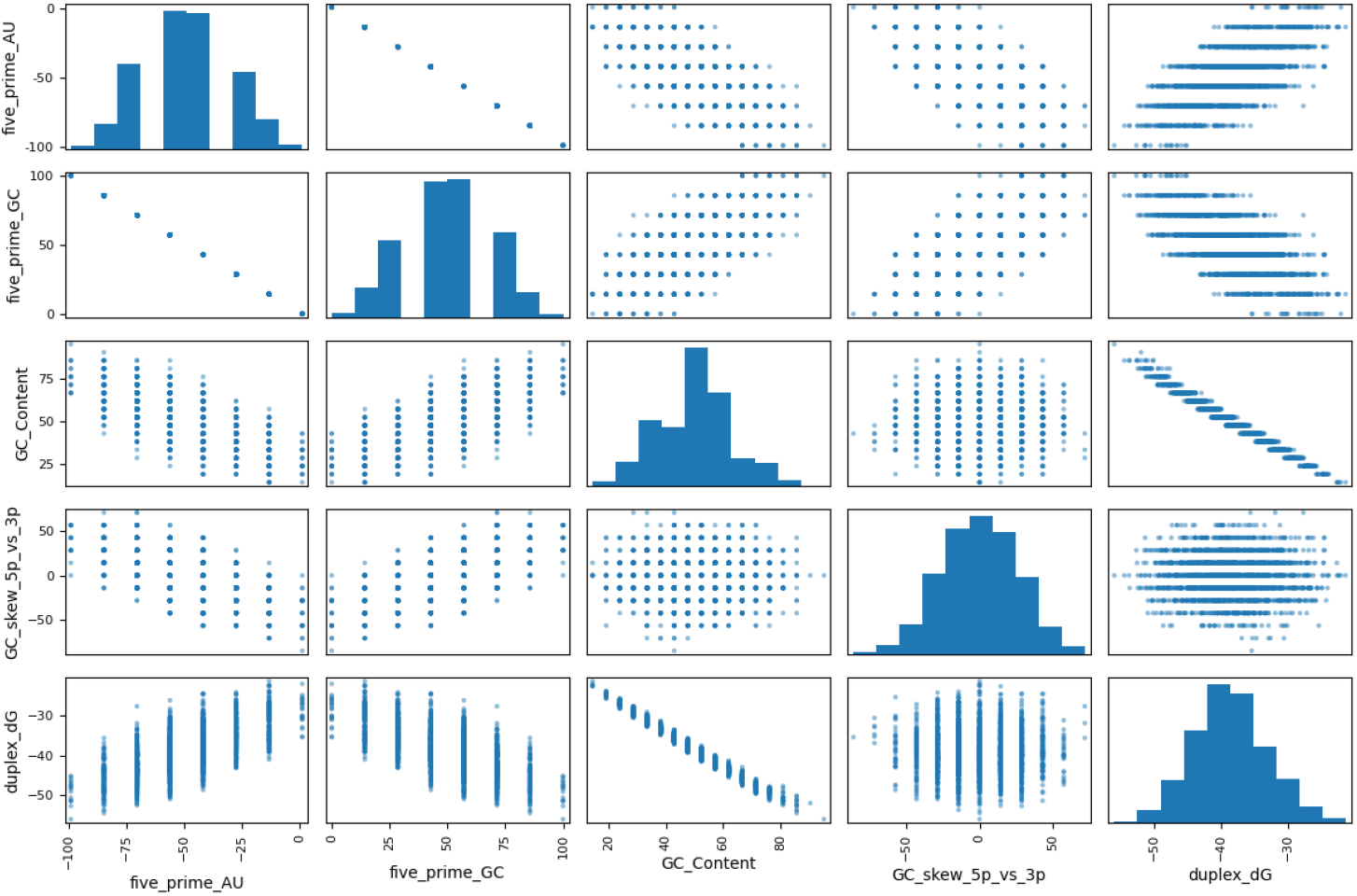
Pairwise scatterplots and marginal distributions for selected thermodynamic and compositional features. Features shown include AU and GC content at the 5^′^end, overall GC content, GC skew between the 5^′^and 3^′^ends, and duplex binding free energy (Δ*G*).

## Supplementary Note 2: Regression Results

**Table SN.2.** Linear Regression Performance. Models trained on different feature group combinations, sorted by *R* Pearson coefficient. Best test *R*^2^ and *R* values are highlighted. C: Composition features; M: Motif and regulatory elements; T: Thermodynamic stability features; S: Structural complexity features.

| Feature Group | Train |  |  |  | Test |  |  |  |
| --- | --- | --- | --- | --- | --- | --- | --- | --- |
| | MAE | RMSE | $R^2$ | $R$ | MAE | RMSE | $R^2$ | $R$ |
| C + M + T + S | 0.112 | 0.141 | 0.516 | 0.718 | **0.112** | **0.144** | **0.454** | **0.676** |
| C + M + S | 0.113 | 0.141 | 0.515 | 0.718 | 0.113 | 0.145 | 0.452 | 0.675 |
| C + M + T | 0.113 | 0.141 | 0.512 | 0.716 | 0.113 | 0.145 | 0.451 | 0.674 |
| C + M | 0.113 | 0.141 | 0.512 | 0.716 | 0.113 | 0.145 | 0.451 | 0.674 |
| C + T + S | 0.113 | 0.141 | 0.513 | 0.716 | 0.113 | 0.145 | 0.451 | 0.674 |
| C + S | 0.113 | 0.141 | 0.513 | 0.716 | 0.113 | 0.145 | 0.449 | 0.673 |
| C + T | 0.113 | 0.141 | 0.510 | 0.714 | 0.114 | 0.145 | 0.448 | 0.672 |
| C | 0.113 | 0.142 | 0.510 | 0.714 | 0.114 | 0.145 | 0.448 | 0.672 |
| M + T | 0.145 | 0.180 | 0.203 | 0.451 | 0.139 | 0.175 | 0.195 | 0.446 |
| M + T + S | 0.144 | 0.180 | 0.209 | 0.458 | 0.139 | 0.175 | 0.195 | 0.446 |
| M + S | 0.145 | 0.182 | 0.194 | 0.440 | 0.142 | 0.177 | 0.175 | 0.423 |
| M | 0.147 | 0.183 | 0.179 | 0.423 | 0.143 | 0.179 | 0.161 | 0.409 |
| T | 0.155 | 0.192 | 0.097 | 0.311 | 0.146 | 0.181 | 0.140 | 0.387 |
| T + S | 0.154 | 0.191 | 0.108 | 0.329 | 0.145 | 0.181 | 0.139 | 0.380 |
| S | 0.158 | 0.196 | 0.062 | 0.248 | 0.151 | 0.188 | 0.074 | 0.275 |

**Table SN.3.** Support Vector Regression. Models trained on different feature group combinations, sorted by *R* Pearson coefficient. Best test *R*^2^ and *R* values are highlighted. C: Composition features; M: Motif and regulatory elements; T: Thermodynamic stability features; S: Structural complexity features.

| Feature Group | Train |  |  |  | Test |  |  |  |
| --- | --- | --- | --- | --- | --- | --- | --- | --- |
| | MAE | RMSE | $R^2$ | $R$ | MAE | RMSE | $R^2$ | $R$ |
| C + M + S | 0.011 | 0.014 | 0.995 | 0.998 | **0.106** | **0.136** | **0.516** | **0.719** |
| C + M + T + S | 0.011 | 0.015 | 0.995 | 0.997 | 0.106 | 0.136 | 0.515 | 0.719 |
| C + M + T | 0.011 | 0.015 | 0.994 | 0.997 | 0.107 | 0.136 | 0.513 | 0.718 |
| C + S | 0.011 | 0.014 | 0.995 | 0.998 | 0.107 | 0.136 | 0.513 | 0.718 |
| C + T + S | 0.011 | 0.014 | 0.995 | 0.998 | 0.107 | 0.136 | 0.513 | 0.717 |
| C + M | 0.011 | 0.014 | 0.995 | 0.997 | 0.107 | 0.136 | 0.513 | 0.717 |
| C + T | 0.011 | 0.014 | 0.995 | 0.997 | 0.108 | 0.137 | 0.508 | 0.715 |
| C | 0.011 | 0.014 | 0.995 | 0.998 | 0.108 | 0.137 | 0.508 | 0.714 |
| M + T | 0.118 | 0.162 | 0.360 | 0.600 | 0.145 | 0.182 | 0.135 | 0.407 |
| T | 0.147 | 0.188 | 0.133 | 0.365 | 0.146 | 0.183 | 0.125 | 0.359 |
| M + S | 0.115 | 0.160 | 0.374 | 0.611 | 0.149 | 0.189 | 0.067 | 0.358 |
| M | 0.135 | 0.175 | 0.255 | 0.505 | 0.148 | 0.185 | 0.098 | 0.351 |
| M + T + S | 0.104 | 0.150 | 0.453 | 0.673 | 0.151 | 0.191 | 0.040 | 0.345 |
| T + S | 0.139 | 0.181 | 0.197 | 0.444 | 0.149 | 0.189 | 0.060 | 0.285 |
| S | 0.153 | 0.192 | 0.093 | 0.306 | 0.152 | 0.190 | 0.054 | 0.245 |

**Table SN.4.** Gradient Boosting. Models trained on different feature group combinations, sorted by *R* Pearson coefficient. Best test *R*^2^ and *R* values are highlighted. C: Composition features; M: Motif and regulatory elements; T: Thermodynamic stability features; S: Structural complexity features.

| Feature Group | Train |  |  |  | Test |  |  |  |
| --- | --- | --- | --- | --- | --- | --- | --- | --- |
| | MAE | RMSE | $R^2$ | $R$ | MAE | RMSE | $R^2$ | $R$ |
| C | 0.0778 | 0.0981 | 0.7643 | 0.8743 | **0.1097** | **0.1418** | **0.4731** | **0.6894** |
| C + T + S | 0.0747 | 0.0940 | 0.7835 | 0.8852 | 0.1109 | 0.1423 | 0.4691 | 0.6863 |
| C + M + T + S | 0.0748 | 0.0943 | 0.7824 | 0.8845 | 0.1115 | 0.1432 | 0.4625 | 0.6809 |
| C + S | 0.0766 | 0.0962 | 0.7735 | 0.8795 | 0.1117 | 0.1438 | 0.4580 | 0.6776 |
| C + M + T | 0.0759 | 0.0956 | 0.7763 | 0.8811 | 0.1120 | 0.1450 | 0.4493 | 0.6709 |
| C + M + S | 0.0760 | 0.0954 | 0.7773 | 0.8817 | 0.1125 | 0.1449 | 0.4497 | 0.6709 |
| C + T | 0.0767 | 0.0965 | 0.7722 | 0.8787 | 0.1121 | 0.1450 | 0.4490 | 0.6704 |
| C + M | 0.0773 | 0.0974 | 0.7680 | 0.8764 | 0.1125 | 0.1454 | 0.4463 | 0.6690 |
| M + T | 0.1138 | 0.1434 | 0.4964 | 0.7045 | 0.1381 | 0.1739 | 0.2074 | 0.4582 |
| M + T + S | 0.1088 | 0.1377 | 0.5355 | 0.7318 | 0.1398 | 0.1764 | 0.1844 | 0.4350 |
| M | 0.1403 | 0.1750 | 0.2500 | 0.5000 | 0.1439 | 0.1806 | 0.1458 | 0.3923 |
| M + S | 0.1213 | 0.1533 | 0.4250 | 0.6519 | 0.1440 | 0.1838 | 0.1148 | 0.3688 |
| T | 0.1247 | 0.1563 | 0.4019 | 0.6340 | 0.1453 | 0.1817 | 0.1349 | 0.3675 |
| T + S | 0.1193 | 0.1500 | 0.4490 | 0.6701 | 0.1455 | 0.1828 | 0.1240 | 0.3535 |
| S | 0.1391 | 0.1740 | 0.2587 | 0.5087 | 0.1536 | 0.1933 | 0.0210 | 0.2093 |

**Fig. SN.2.**
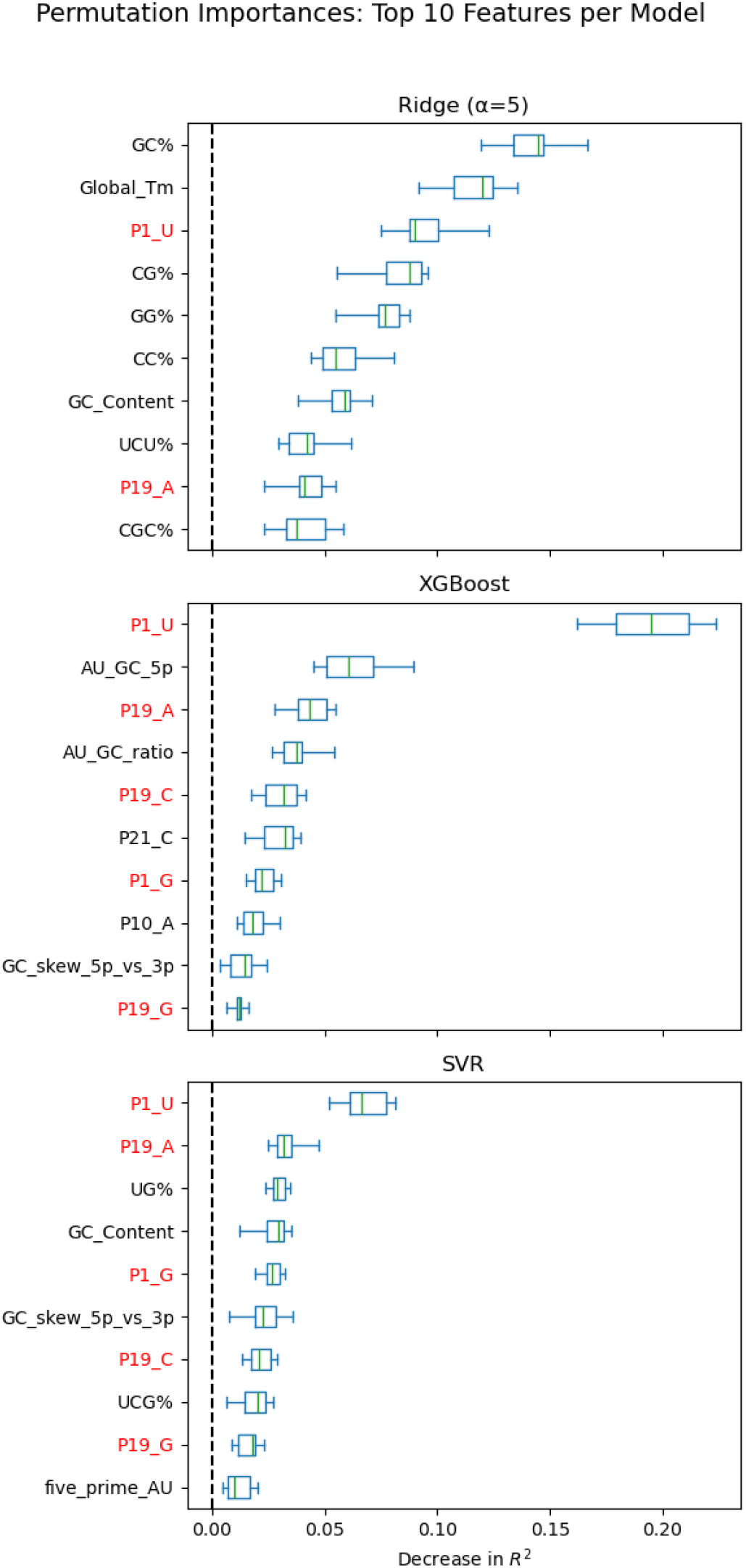
Permutation importances for the top 10 features from three regression models (Ridge regression with *α* = 5, XGBoost, and SVR), each trained on trained on its best-performing feature group as determined by test Pearson correlation. Feature importance was measured as the mean decrease in *R*^2^ across 10 permutations, with with box plots representing the distribution across repeats. Position-specific nucleotides (e.g., P1_U, P19_A) consistently ranked among the most important features across models, supporting the robustness of the SVR results presented in the main text (Figure 1).

## Supplementary Note 3: Classification Results

**Table SN.5.** Logistic regression. Comparison of models trained on different feature group combinations using two label thresholds (0.5 and 0.7), sorted by Test ROC. C: Composition; M: Motifs and regulatory elements; T: Thermodynamic stability; S: Structural complexity.

| Feature Group | Train |  |  | Test |  |  |
| --- | --- | --- | --- | --- | --- | --- |
|  | Accuracy | F1 | ROC | Accuracy | F1 | ROC |
| *Label threshold: 0.5* |  |  |  |  |  |  |
| C + M + T | 0.859 | 0.858 | 0.931 | **0.801** | **0.809** | **0.885** |
| C + M | 0.859 | 0.858 | 0.931 | 0.801 | 0.809 | 0.885 |
| C + M + T + S | 0.861 | 0.860 | 0.933 | 0.795 | 0.809 | 0.882 |
| C + M + S | 0.860 | 0.859 | 0.933 | 0.795 | 0.809 | 0.882 |
| C | 0.852 | 0.851 | 0.927 | 0.795 | 0.802 | 0.870 |
| C + T | 0.853 | 0.853 | 0.928 | 0.807 | 0.811 | 0.868 |
| C + T + S | 0.859 | 0.859 | 0.929 | 0.795 | 0.800 | 0.865 |
| C + S | 0.858 | 0.858 | 0.929 | 0.795 | 0.800 | 0.864 |
| M + T + S | 0.658 | 0.662 | 0.725 | 0.673 | 0.678 | 0.721 |
| M + T | 0.656 | 0.657 | 0.720 | 0.637 | 0.640 | 0.717 |
| M + S | 0.650 | 0.652 | 0.713 | 0.667 | 0.671 | 0.707 |
| M | 0.642 | 0.644 | 0.696 | 0.643 | 0.647 | 0.703 |
| T | 0.586 | 0.615 | 0.634 | 0.579 | 0.613 | 0.636 |
| T + S | 0.604 | 0.643 | 0.638 | 0.573 | 0.614 | 0.630 |
| S | 0.578 | 0.618 | 0.599 | 0.503 | 0.581 | 0.588 |
| *Label threshold: 0.7* |  |  |  |  |  |  |
| C + M + T + S | 0.804 | 0.804 | 0.888 | **0.740** | **0.740** | **0.839** |
| C + M + S | 0.804 | 0.804 | 0.888 | 0.744 | 0.745 | 0.838 |
| C + M + T | 0.805 | 0.805 | 0.886 | 0.740 | 0.742 | 0.836 |
| C + M | 0.804 | 0.805 | 0.886 | 0.738 | 0.741 | 0.836 |
| C + T + S | 0.802 | 0.803 | 0.886 | 0.729 | 0.728 | 0.836 |
| C + S | 0.803 | 0.803 | 0.886 | 0.733 | 0.733 | 0.835 |
| C + T | 0.801 | 0.801 | 0.884 | 0.735 | 0.737 | 0.833 |
| C | 0.801 | 0.801 | 0.884 | 0.731 | 0.733 | 0.833 |
| M + T + S | 0.662 | 0.669 | 0.733 | 0.662 | 0.662 | 0.731 |
| M + T | 0.661 | 0.665 | 0.729 | 0.654 | 0.654 | 0.722 |
| M + S | 0.654 | 0.660 | 0.721 | 0.673 | 0.671 | 0.719 |
| M | 0.654 | 0.648 | 0.715 | 0.648 | 0.629 | 0.707 |
| T + S | 0.612 | 0.636 | 0.661 | 0.637 | 0.669 | 0.665 |
| T | 0.607 | 0.621 | 0.656 | 0.610 | 0.628 | 0.653 |
| S | 0.564 | 0.608 | 0.598 | 0.573 | 0.611 | 0.615 |

**Table SN.6.** Support Vector Machine (SVM) classification. Comparison of models trained on different feature group combinations under two label thresholds (0.5 and 0.7), sorted by Test ROC. C: Composition; M: Motifs and regulatory elements; T: Thermodynamic stability; S: Structural complexity.

| Feature Group | Train |  |  | Test |  |  |
| --- | --- | --- | --- | --- | --- | --- |
|  | Accuracy | F1 | ROC | Accuracy | F1 | ROC |
| *Label threshold: 0.5* |  |  |  |  |  |  |
| C + M | 0.937 | 0.937 | 0.984 | **0.708** | **0.713** | **0.806** |
| C + M + T | 0.936 | 0.936 | 0.983 | 0.719 | 0.724 | 0.805 |
| C + M + S | 0.933 | 0.933 | 0.983 | 0.702 | 0.705 | 0.801 |
| C | 0.938 | 0.937 | 0.987 | 0.713 | 0.713 | 0.800 |
| C + M + T + S | 0.932 | 0.932 | 0.982 | 0.702 | 0.702 | 0.800 |
| C + T | 0.937 | 0.937 | 0.986 | 0.702 | 0.702 | 0.799 |
| C + T + S | 0.938 | 0.938 | 0.985 | 0.702 | 0.702 | 0.794 |
| C + S | 0.941 | 0.940 | 0.986 | 0.713 | 0.713 | 0.794 |
| M + T | 0.714 | 0.713 | 0.794 | 0.643 | 0.647 | 0.681 |
| M + T + S | 0.734 | 0.731 | 0.814 | 0.614 | 0.593 | 0.676 |
| M | 0.679 | 0.640 | 0.741 | 0.602 | 0.547 | 0.674 |
| M + S | 0.720 | 0.711 | 0.793 | 0.585 | 0.548 | 0.660 |
| T | 0.617 | 0.660 | 0.662 | 0.550 | 0.613 | 0.613 |
| S | 0.603 | 0.677 | 0.645 | 0.550 | 0.645 | 0.588 |
| T + S | 0.648 | 0.672 | 0.702 | 0.544 | 0.571 | 0.575 |
| *Label threshold: 0.7* |  |  |  |  |  |  |
| C + S | 0.928 | 0.928 | 0.983 | **0.760** | **0.763** | **0.844** |
| C | 0.926 | 0.926 | 0.982 | 0.754 | 0.759 | 0.844 |
| C + T | 0.923 | 0.923 | 0.981 | 0.754 | 0.759 | 0.843 |
| C + T + S | 0.925 | 0.925 | 0.981 | 0.758 | 0.761 | 0.843 |
| C + M | 0.921 | 0.921 | 0.980 | 0.754 | 0.756 | 0.840 |
| C + M + S | 0.925 | 0.925 | 0.980 | 0.758 | 0.761 | 0.840 |
| C + M + T + S | 0.921 | 0.921 | 0.979 | 0.758 | 0.761 | 0.839 |
| C + M + T | 0.918 | 0.918 | 0.979 | 0.760 | 0.763 | 0.839 |
| M + T | 0.695 | 0.696 | 0.767 | 0.658 | 0.651 | 0.708 |
| M + T + S | 0.714 | 0.720 | 0.793 | 0.652 | 0.651 | 0.707 |
| M | 0.679 | 0.680 | 0.736 | 0.642 | 0.632 | 0.693 |
| M + S | 0.709 | 0.717 | 0.776 | 0.642 | 0.648 | 0.690 |
| T + S | 0.637 | 0.669 | 0.685 | 0.625 | 0.667 | 0.659 |
| T | 0.613 | 0.637 | 0.660 | 0.621 | 0.649 | 0.653 |
| S | 0.593 | 0.651 | 0.623 | 0.621 | 0.670 | 0.617 |

**Table SN.7.** Random Forest. Models trained on different group combinations, sorted by test ROC. Rows are grouped by label threshold (0.5 or 0.7). C: Composition; M: Motifs and regulatory elements; T: Thermodynamic stability; S: Structural complexity.

| Feature Group | Train |  |  | Test |  |  |
| --- | --- | --- | --- | --- | --- | --- |
|  | Accuracy | F1 | ROC | Accuracy | F1 | ROC |
| *Label threshold: 0.5* |  |  |  |  |  |  |
| C + T | 0.694 | 0.685 | 0.755 | **0.784** | **0.784** | **0.852** |
| C | 0.702 | 0.688 | 0.763 | 0.719 | 0.711 | 0.828 |
| C + S | 0.706 | 0.692 | 0.760 | 0.743 | 0.732 | 0.819 |
| C + M + T | 0.693 | 0.683 | 0.767 | 0.754 | 0.750 | 0.816 |
| C + M | 0.702 | 0.685 | 0.756 | 0.743 | 0.732 | 0.811 |
| C + M + S | 0.705 | 0.692 | 0.753 | 0.696 | 0.683 | 0.808 |
| C + M + T + S | 0.712 | 0.700 | 0.760 | 0.719 | 0.718 | 0.792 |
| C + T + S | 0.693 | 0.681 | 0.765 | 0.713 | 0.713 | 0.789 |
| M + S | 0.596 | 0.570 | 0.629 | 0.626 | 0.628 | 0.693 |
| M + T + S | 0.615 | 0.602 | 0.653 | 0.643 | 0.647 | 0.687 |
| M + T | 0.617 | 0.604 | 0.660 | 0.632 | 0.644 | 0.669 |
| S | 0.502 | 0.484 | 0.507 | 0.620 | 0.620 | 0.638 |
| M | 0.614 | 0.618 | 0.642 | 0.602 | 0.585 | 0.618 |
| T | 0.577 | 0.573 | 0.618 | 0.532 | 0.545 | 0.574 |
| T + S | 0.573 | 0.574 | 0.604 | 0.532 | 0.551 | 0.571 |
| *Label threshold: 0.7* |  |  |  |  |  |  |
| C + M | 0.712 | 0.704 | 0.793 | **0.765** | **0.761** | **0.822** |
| C + S | 0.717 | 0.708 | 0.796 | 0.731 | 0.723 | 0.816 |
| C + T + S | 0.730 | 0.724 | 0.796 | 0.738 | 0.731 | 0.811 |
| C | 0.719 | 0.710 | 0.796 | 0.715 | 0.709 | 0.810 |
| C + M + T + S | 0.711 | 0.705 | 0.784 | 0.742 | 0.738 | 0.806 |
| C + M + T | 0.728 | 0.719 | 0.793 | 0.719 | 0.711 | 0.802 |
| C + M + S | 0.713 | 0.703 | 0.791 | 0.735 | 0.727 | 0.801 |
| C + T | 0.714 | 0.705 | 0.794 | 0.727 | 0.725 | 0.800 |
| M + T + S | 0.629 | 0.629 | 0.683 | 0.673 | 0.671 | 0.719 |
| M + T | 0.623 | 0.620 | 0.674 | 0.681 | 0.678 | 0.706 |
| M + S | 0.610 | 0.611 | 0.663 | 0.635 | 0.633 | 0.698 |
| M | 0.620 | 0.612 | 0.678 | 0.631 | 0.621 | 0.675 |
| T + S | 0.589 | 0.595 | 0.638 | 0.600 | 0.608 | 0.665 |
| T | 0.582 | 0.583 | 0.623 | 0.615 | 0.617 | 0.654 |
| S | 0.565 | 0.573 | 0.575 | 0.562 | 0.568 | 0.589 |

**Fig. SN.3.**
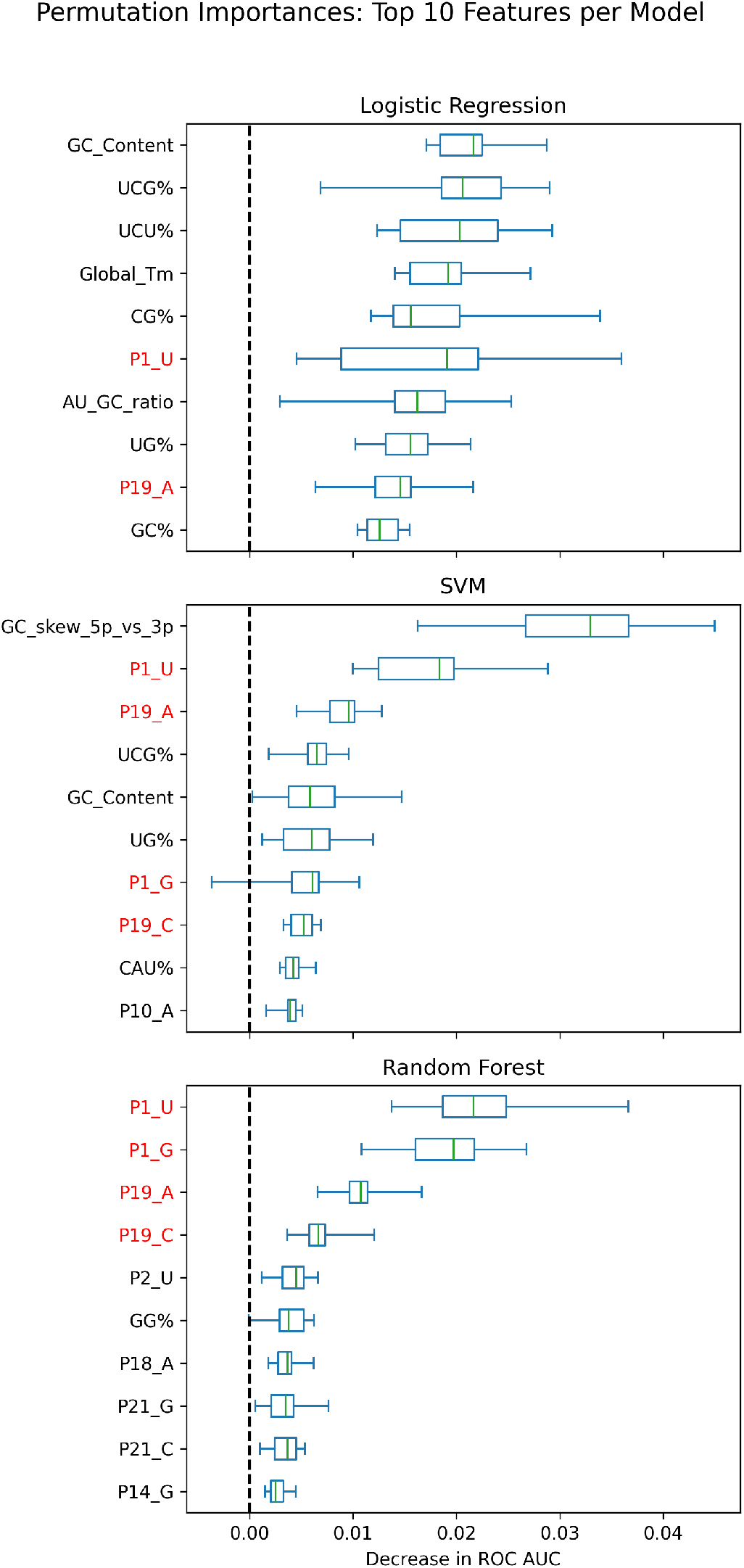
Permutation importances for the top 10 features from three classification models (Logistic Regression, SVM, and Random Forest), each trained on its best-performing feature group as determined by test ROC AUC. Feature importance was measured as the mean decrease in ROC AUC across 10 permutations, with box plots representing the distribution across repeats. Position-specific nucleotides (especially P1_U and P19_A) were among the top predictors for SVM and Random Forest, consistent with regression results. The relative ranking of features varied across models, reflecting differences in how each algorithm captures feature interactions.

## Footnotes

1 While permutation-based feature importance provides a model-agnostic and interpretable approach for quantifying the contribution of each input variable, its application to ordinary least squares (OLS) linear regression proved unstable in our setting. Specifically, the absence of regularization in OLS makes the model highly sensitive to collinearity and small perturbations in the feature space. As permutation importance involves randomly shuffling feature values, this instability led to inflated and erratic importance scores when using unregularized linear regression. To address this, we replaced OLS with Ridge regression (L2-regularized linear regression), which introduces a penalty on large coefficients and is known to be more robust to multicollinearity. We evaluated multiple values of the regularization parameter *α*, and found that Ridge regression with *α* = 5 consistently produced stable results across cross-validation folds and yielded interpretable permutation importance profiles. We therefore report results using Ridge regression throughout.

